# A spectacular anomaly in the 4-mer composition of the giant pandoravirus genomes reveals a stringent new evolutionary selection process

**DOI:** 10.1101/712018

**Authors:** Olivier Poirot, Sandra Jeudy, Chantal Abergel, Jean-Michel Claverie

## Abstract

The Pandoraviridae is a rapidly growing family of giant viruses, all of which have been isolated using laboratory strains of Acanthamoeba. The genomes of ten distinct strains have been fully characterized, reaching up to 2.5 Mb in size. These double-stranded DNA genomes encode the largest of all known viral proteomes and are propagated in oblate virions that are among the largest ever-described (1.2 µm long and 0.5 µm wide). The evolutionary origin of these atypical viruses is the object of numerous speculations. Applying the Chaos Game Representation to the pandoravirus genome sequences, we discovered that the tetranucleotide (4-mer) “AGCT” is totally absent from the genomes of 2 strains (*P. dulcis* and *P. quercus*) and strongly underrepresented in others. Given the amazingly low probability of such an observation in the corresponding randomized sequences, we investigated its biological significance through a comprehensive study of the 4-mer compositions of all viral genomes. Our results indicate that “AGCT” was specifically eliminated during the evolution of the Pandoraviridae and that none of the previously proposed host-virus antagonistic relationships could explain this phenomenon. Unlike the three other families of giant viruses (Mimiviridae, Pithoviridae, Molliviridae) infecting the same Acanthamoeba host, the pandoraviruses exhibit a puzzling genomic anomaly suggesting a highly specific DNA editing in response to a new kind of strong evolutionary pressure.

**Importance:** The recent years have seen the discovery of several families of giant DNA viruses all infecting the ubiquitous amoebozoa of the genus Acanthamoeba. With dsDNA genomes reaching 2.5 Mb in length packaged in oblate particles the size of a bacterium, the pandoraviruses are the most complex and largest viruses known as of today. In addition to their spectacular dimensions, the pandoraviruses encode the largest proportion of proteins without homolog in other organisms, thought to result from a *de novo* gene creation process. While using comparative genomics to investigate the evolutionary forces responsible for the emergence of such an unusual giant virus family, we discovered a unique bias in the tetranucleotide composition of the pandoravirus genomes that can only result from an undescribed evolutionary process not encountered in any other microorganism.

## Introduction

The Pandoraviruses are among the growing number of families of environmental giant DNA viruses infecting protozoans and isolated using the laboratory host Acanthamoeba (Protozoa/Lobosa/Ameobida/ Acanthamoebidae/ Acanthamoeba) ^1–4^. As of today, they exhibit the largest fully characterized viral genomes, made of linear dsDNA molecules from 1.9 to 2.5 Mb in size, predicted to encode up to 2500 proteins^1–3^. After their internalization by phagocytosis, these viruses multiply in their amoebal host through a lytic cycle lasting about 12 hours, ending with the production of hundreds of giant amphora-shaped particles (1.2 µm long and 0.5 µm wide)^1–3^. The phylogenetic structure of the Pandoraviridae family exhibits two separate clusters referred to as A- and B-clades^2,3^ (Fig. 1). Despite this clear phylogenetic signal (computed using a core set of 455 orthologous proteins), strains belonging to clade A or B did not exhibit noticeable differences in terms of virion morphology, infectious cycle, host range, or global genome structure and statistics^1–3^.

**Figure 1.**
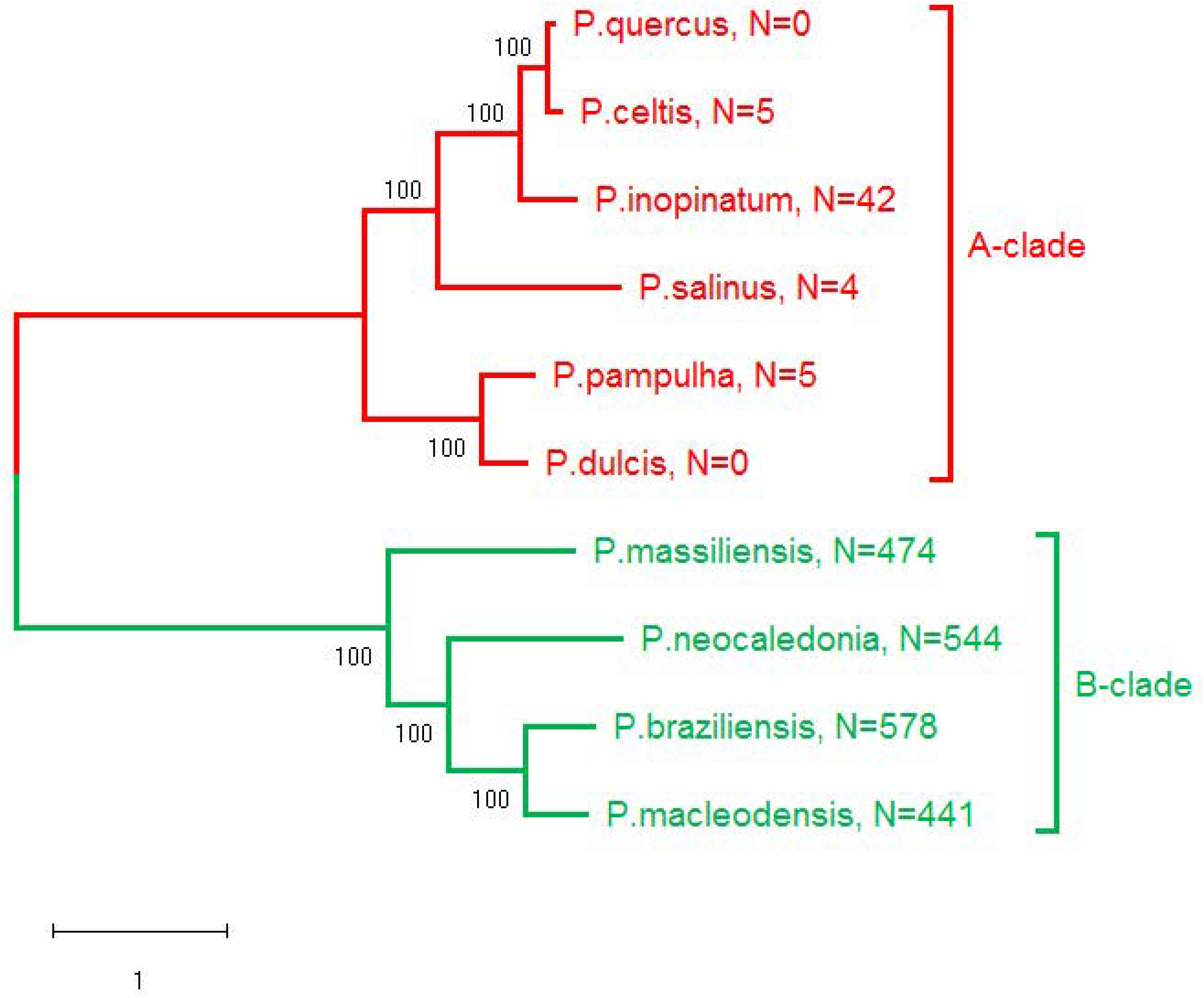
Phylogenetic structure of the Pandoraviridae. Adapted from [ref. 3]. The number of occurrences of the “AGCT” 4-mer is indicated for the genome of each strain. The counts are given for one DNA strand and are identical for both strands (“AGCT” is palindromic).

In addition to their unusual virion morphology and gigantic genomes, the pandoraviruses exhibits other unique features such as an unmatched proportion (>90%) of genes coding for proteins without any database homologs (ORFans) outside of the Pandoraviridae family, and strain-specific genes contributing to an unlimited pan-genome^1–3^. These features, confirmed by the analysis of additional strains^5^, led us to suggest that a process of *de novo* and *in situ* gene creation might be at work in pandoraviruses^2, 3^. Following this history of unexpected findings, we thought that further analyses of the Pandoraviridae might reveal additional surprises.

While searching for hidden genomic patterns eventually linked to evolutionary processes unique to the pandoraviruses, we used a Chaos Game graphical representation of their genome sequences^6–7^. This method converts long one-dimensional DNA sequence into a fractal-like image, through which a human observer may detect specific patterns. This representation illustrates in a holistic manner the frequencies of all oligonucleotides of arbitrary length k (k-mers) in a given DNA sequence. Using this approach led us to discover that the 4-mer “AGCT” was uniquely absent from the genome of *Pandoravirus dulcis*, providing the starting point of the present study.

## Results

### The absence of any given 4-mer in a long random DNA sequence is highly improbable

After detecting the absence of the “AGCT” word in the Chaos Game graphical representation of the *P. dulcis* genome, we computed the number of occurrence of all 4-mers in the ten available Pandoravirus genome sequences using direct counting^8^. This revealed that “AGCT” was also absent from the genome of *P. quercus*. Notice that although these strains belong to the same A-clade, their genome sequences are nevertheless far from identical (their orthologous proteins share 72% identical residues in average), hence the common missing “AGCT” is not a mere consequence of their sequence similarity.

Such a plain finding might not sound very interesting, until one realize that not encountering a single occurrence of “AGCT” in DNA sequences respectively 1.908.524 bp (*P. dulcis*) and 2.077.288 bp (*P. quercus*) is amazingly unlikely, as shown below, using increasingly sophisticated computations.

In the simplest case, let us first consider a random DNA sequence with equal proportions of the four nucleotides (%A=%T=%C=%G=25%). Since there are 256 distinct 4-mers, the probability for each of them to occur at a given position in an increasingly long sequence tends to *p*_AGCT_ = 1/_256_. In a random sequence of approximately 2 Mbp, one thus expects an average of about 7800 occurrences for each distinct 4-mers. This already suggests how unlikely it is for one of them to be absent.

To estimate the order of magnitude of such probability, the DNA sequence is seen as consisting of 4 sets of non-overlapping 4-mers collected according to 4 different “reading frames” (e.g. 4-mers 1-4, 5-8, 9-12, …, etc, for frame 1). The different reading frames thus correspond to approximately 500,000 positions each.

At each of these position, the probability for “AGCT” not to occur is *q*_AGCT_ = 255/_256_. For one reading frame, this probability becomes approximately

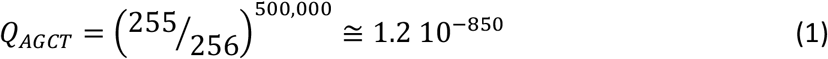

and:

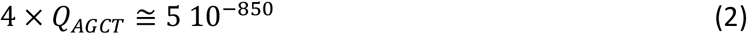

for the 4 reading frames (assuming them to be independent for the sake of simplicity).

Such a value is smaller than any that could be computed in reference to a physical process. For instance, one second approximately corresponds to 2 10^−18^ of the age of the universe.

The above probability should actually be corrected to account for the fact that we did not specifically search for “AGCT” while analyzing the viral genome. Any missing 4-mer would have raised the same interest. A Bonferroni correction should then be applied to compensate for the multiple testing of 256 different 4-mers. However, the probability of not finding any 4-mer, *Q*_*any*_, remains an incommensurably small number.

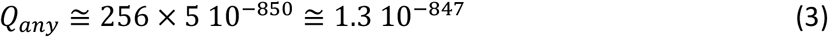

We may further argue that this event was bound to occur in at least one genome given the huge amount of DNA sequence that is now available, for instance in Genbank. The calculation runs as follows; The april 2019 release of Genbank contains about 3.2 10^11^bp. Assuming that all Genbank entries are 2 Mb-long sequences, this would correspond to 1.6 10^5^ theoretical pandoravirus genomes. The order of magnitude of the probability of observing one of them missing any of the 4-mers remains amazingly small at about

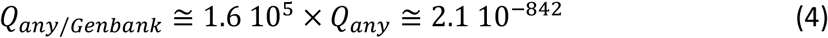

Finally, one may want to make a final adjustment by taking into account that the *P. dulcis* genome is 64% G+C rich. This slightly change the probability of random occurrence of “AGCT” from p_AGCT_ = 1/_256_ = 0.00391 to

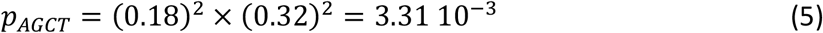

then

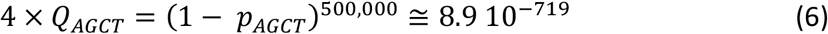

Using the same Bonferroni correction as above lead to the final conservative estimate:

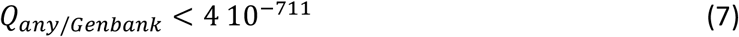

still an incommensurably small probability (e.g. the same as not getting a single head in 2360 tosses of a fair coin).

As the above computation remains an approximation (neglecting the overlap of neighboring 4-mers), we estimated how unlikely it is that any 4-mer would be missing from large DNA sequences by a different approach. We computer generated a large number of random sequences of increasing sizes and recorded the threshold at which point none of the 4-mers is missing. Fig. 2 displays the results of such computer experiment. It shows how fast the probability of any 4-mer missing is decreasing with the random sequence size. In this experiment, we found that the proportion of sequences larger than 10,000 bp missing anyone of the 256 4-mers was less than 1/10,000.

**Figure 2.**
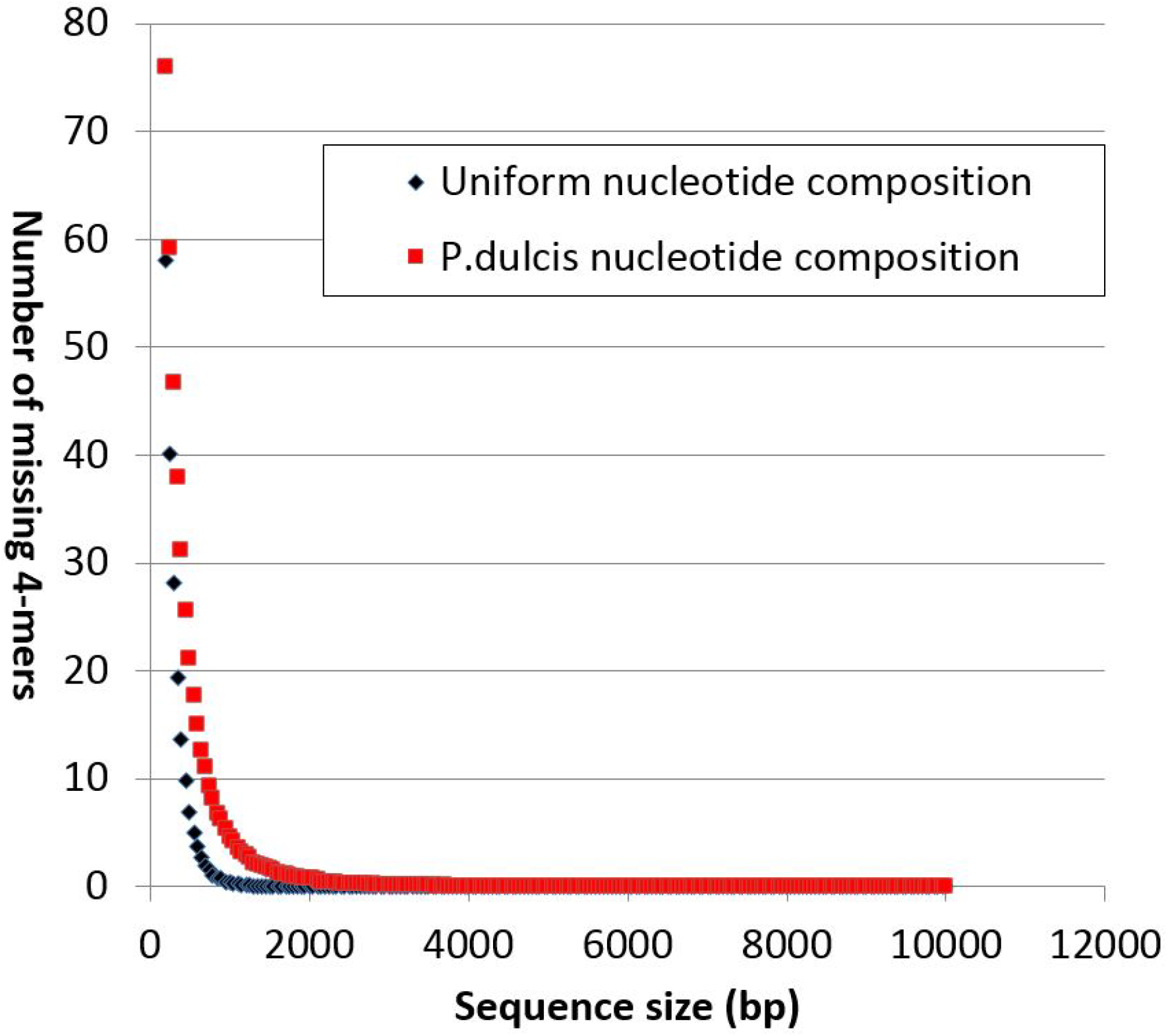
Influence of random sequence length on the number of missing 4-mers. 10.000 random sequences up to 10.000 bp in size were analyzed. Except for extremely rare fluctuations, no sequence longer than 4000 bp exhibits a missing 4-mer. 4-mer overlaps as well as nucleotide compositions are taken into account in this analysis.

### Caveat: randomized sequences exhibit strongly unnatural 4-mer distributions

The above sections already suggested that it is impossible for the *P. dulcis* and *P. quercus* genomes to be missing “AGCT” solely by chance without invoking a biological constraint. However, this conclusion rests on the assumption that the randomization process suitably modeled these genomes. However, a comparison of the frequency distribution of the various 4-mers found in the actual *P. dulcis* genome (and of other pandoraviruses) with that in its randomized sequence shows spectacular differences (Fig. 3). While the natural sequence consist of 4-mers occurring at frequencies distributed along a large and rather continuous interval, the randomized sequence exhibits 4-mers occurring around 5 narrow peaks of frequencies with none in between. As expected from a good quality randomization, these peaks corresponds to the frequencies of the five types of 4-mers: those consisting of only A or T at the lower end, those consisting of only G or C at the higher end, and those consisting of (A or T)/(G or C) in proportions 1/3, 2/2, and 3,1 in between. The more continuous and spread out natural distribution is the testimony of multiple evolutionary constraints, most of them unknown, that have resulted in a distinct 4-mer usage, like a dialect or a language tic inherited from past generations^9^.

**Figure 3.**
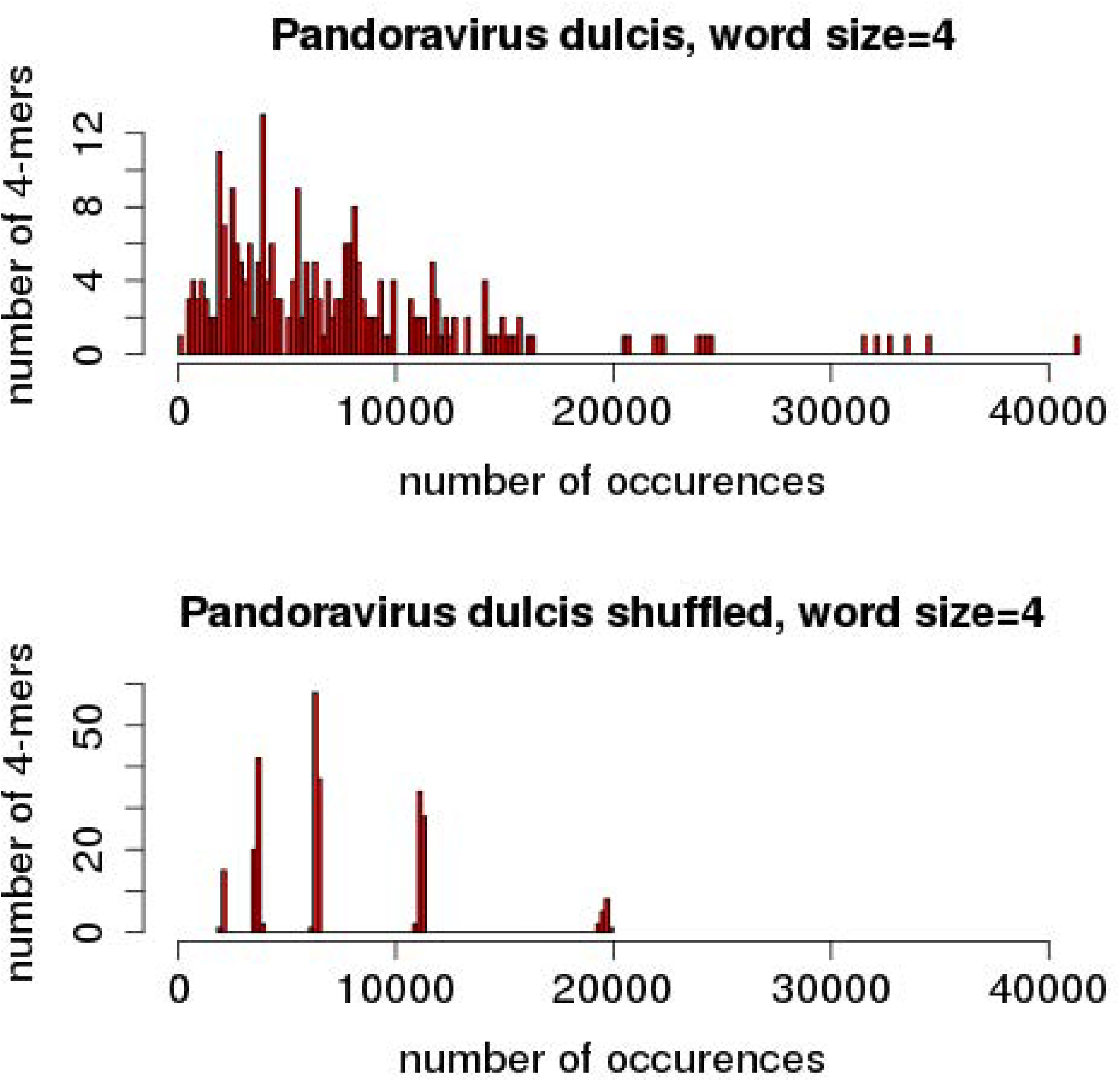
Distribution of 4-mer frequencies in natural and randomized genome sequences. Top: histogram of the number of distinct 4-mers occurring at various numbers of occurrences in the *P. dulcis* genome; Bottom: same analysis after randomization.

First, notice that the missing “AGCT” does not correspond to the 4-mer type with the lowest expected frequency (but the middle one). Second, it is clear that the above probability calculations based on such distorted model of the natural sequence, cannot be used as a reliable estimate of statistical significance. This problem is similar to the one encountered when trying to evaluate the quality of local sequence alignments in similarity searches^10, 11^.

We can mitigate the effect of the above stringent randomization (only preserving the original nucleotide composition) by using the *P. dulcis* and *P. quercus* actual genome sequences to evaluate to what extent the absence of “AGCT” might be the mere statistical consequence of the frequency of its constituent 3-mers: AGC and GCT.

As shown in Table 1, AGC and GCT are not among the least frequent 3-mers found in the *P. dulcis* or *P. quercus* genomes. As the theoretical average is 1/64 (≈ 0.0156), their proportions range from 0.0156 to 0.0097 within the coding and non-coding regions of the genomes. On one given strand, AGC and GCT also do not strongly segregate from each other’s in coding *versus* intergenic regions (Table 1). By combining the AGC 3-mer frequency with that of the single nucleotide T (p _(t)_ =0.182 for *P. dulcis*, p _(t)_ =0.196 for *P. quercus*), the expected number of “AGCT” per strand is 4286 for *P. dulcis* and 4898 for *P. quercus*, while none is observed. Such stark contrast between expected and observed values is unique to the “AGCT” 4-mer. By comparison, the palindromic “ACGT” 4-mer (with an identical composition) exhibits a statistical behavior (Table 1, grey bottom lines) much closer to the 3-mer-dependent random sequence model.

**Table 1.**
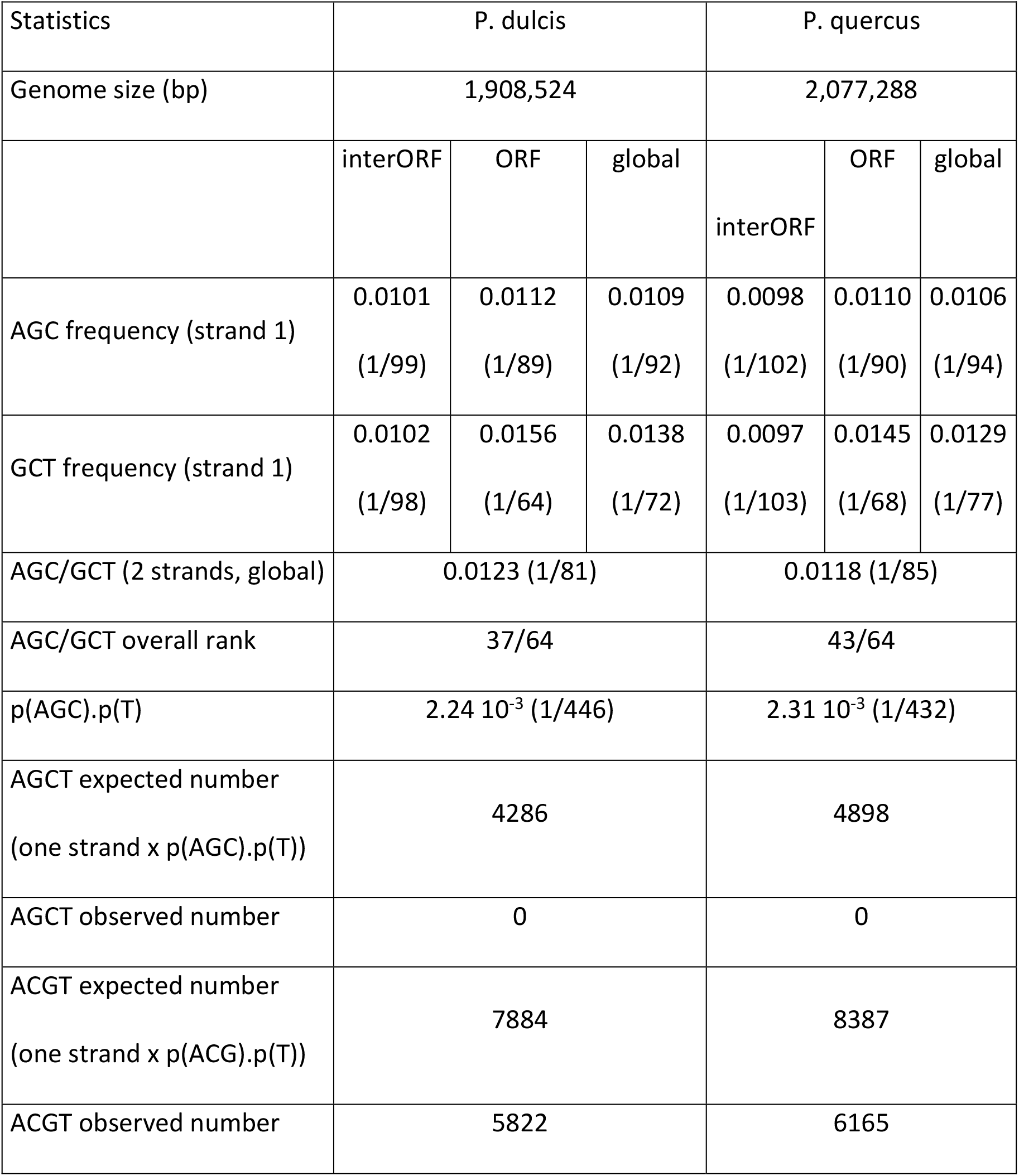
Distribution of the AGC (and the complementary GCT) 3-mers

| Statistics | P. dulcis |  |  | P. quercus |  |  |
| --- | --- | --- | --- | --- | --- | --- |
| Genome size (bp) | 1,908,524 |  |  | 2,077,288 |  |  |
|  | interORF | ORF | global | interORF | ORF | global |
| AGC frequency (strand 1) | 0.0101<br>(1/99) | 0.0112<br>(1/89) | 0.0109<br>(1/92) | 0.0098<br>(1/102) | 0.0110<br>(1/90) | 0.0106<br>(1/94) |
| GCT frequency (strand 1) | 0.0102<br>(1/98) | 0.0156<br>(1/64) | 0.0138<br>(1/72) | 0.0097<br>(1/103) | 0.0145<br>(1/68) | 0.0129<br>(1/77) |
| AGC/GCT (2 strands, global) | 0.0123 (1/81) |  |  | 0.0118 (1/85) |  |  |
| AGC/GCT overall rank | 37/64 |  |  | 43/64 |  |  |
| p(AGC).p(T) | $2.24 \cdot 10^{-3}$ (1/446) | | | $2.31 \cdot 10^{-3}$ (1/432) | | |
| AGCT expected number<br>(one strand x p(AGC).p(T)) | 4286 |  |  | 4898 |  |  |
| AGCT observed number | 0 |  |  | 0 |  |  |
| ACGT expected number<br>(one strand x p(ACG).p(T)) | 7884 |  |  | 8387 |  |  |
| ACGT observed number | 5822 |  |  | 6165 |  |  |

### No 4-mer is missing from the largest actual viral genomes

As vividly illustrated in Fig. 3, the 4-mer distributions in randomized sequences strongly depart from that in natural genomes. We thus analyzed all complete genome sequences available in the viral section of Genbank^12^, to investigate to what extent the absence of a given 4-mer was exceptional for genomes in the size range corresponding to Pandoraviruses.

We found that the next largest viral genomes missing a 4-mers were those of five phages infecting enterobacteria, with unusual genome sizes in the 345kb-359kb range^13–16^. Except for *P. dulcis* and *P. quercus*, none of the 26 largest publicly available viral genomes (including 25 large/giant eukaryotic viruses, and phage G) ^12^ were missing a 4-mer (Fig. 4). Thus, even by comparison with natural sequences, *P. dulcis* and *P. quercus* appear truly exceptional.

**Figure 4.**
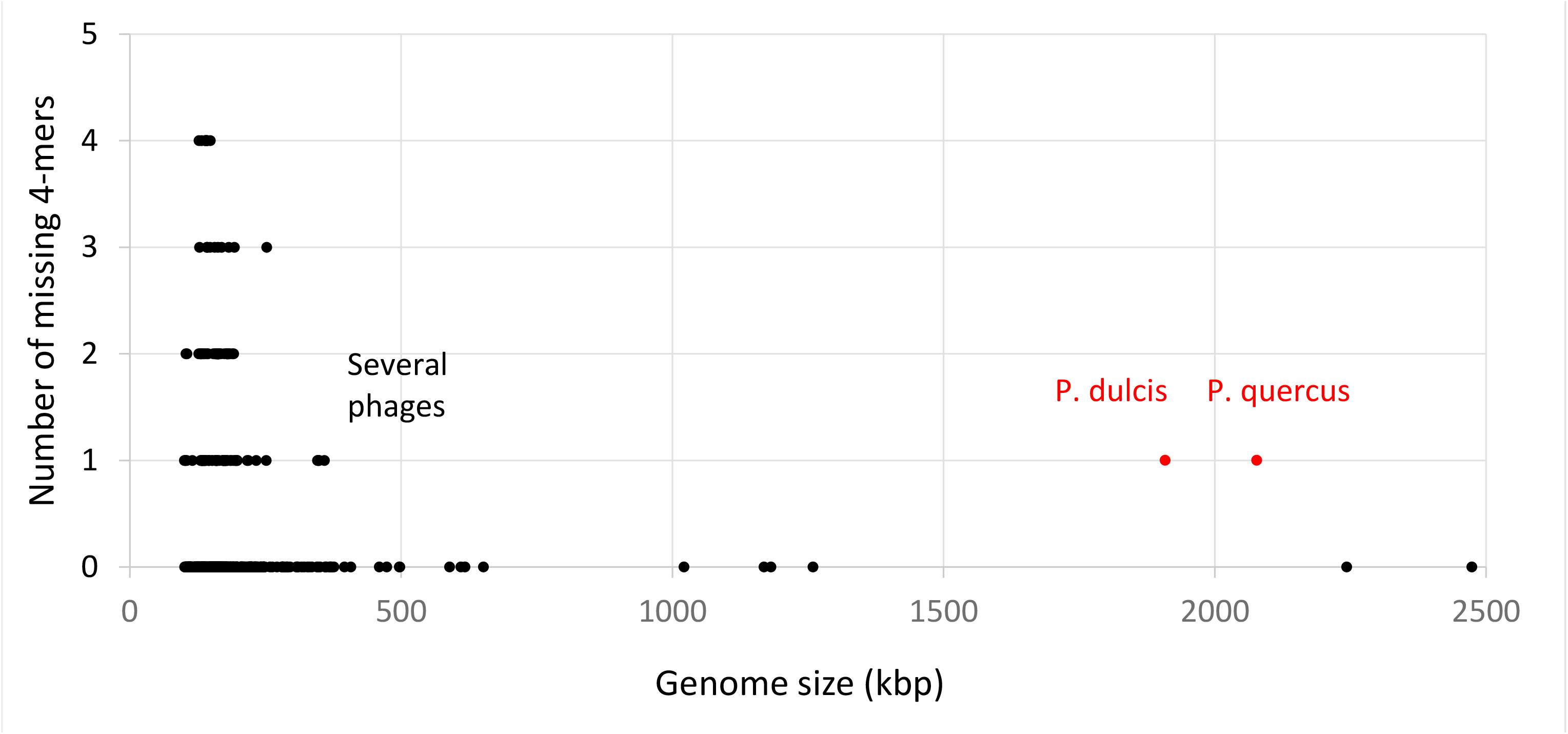
Missing 4-mers in the largest viral genomes. Except for *P. dulcis* and *P. quercus*, the largest viral genomes missing a 4-mers are those of 5 distinct bacteriophages (accession numbers: NC_019401, NC_025447, NC_027364, NC_027399, NC_019526).

We noticed that the five large enterobacteria-infecting phages pointed out by our analysis, were all missing the same “GCGC” 4_mer although they exhibit divergent genomic sequences and were isolated from different hosts^13–16^. This palindromic 4-mer might be the target of isoschizomeric restriction endonucleases functionally homologous to HhaI found in *Haemophilus haemolyticus*, a Gammaproteobacteria. Many of them have been described (see https://enzymefinder.neb.com). We will return to the hypothesis that some 4-mers might be missing in response to a host or viral defense mechanism^17^ in the discussion section.

### The anomalous distribution of “AGCT” correlates with the Pandoraviridae phylogenetic structure

The absence of “AGCT” in *P. dulcis* and *P. quercus* genomes becomes even more intriguing when put in the context of the phylogenetic structure of the whole pandoravirus family. As shown in Fig. 1, the Pandoraviridae neatly cluster into two separate clades. For well-conserved proteins (such as the DNA polB), the percentage of identical residues between intra-clade orthologs is in the 82% to 90% range, and in the 72% to 76% range between the two clades. The corresponding genome sequences are thus far from being identical (and only partially collinear) within each clade. It is thus quite remarkable that the “AGCT” count exhibits a consistent trends to be very low in A-clade members, and at least 10 times higher in B-clade strains. Such a contrast was strong enough to pre-classify three unpublished isolates prior complete genome assembly and finishing (data not shown).

The large difference in “AGCT” counts could be eventually due to the deletion of a genomic region concentrating most of them, for instance within a repeated structure absent from the A-clade isolates. However, Fig. 5 shows that this is not at all the case. In B-clade isolates, the numerous occurrences of “AGCT” are rather uniformly distributed along the whole genomes. However, we noticed that the “AGCT” distribution in the *P. neocaledonia* genome exhibits a change of slope at one of its extremity, as if the corresponding segment had been acquired from a A-clade strain. Such hypothesis was confirmed using a dot-plot comparison with the *P. salinus* genome, to which this terminal segment is clearly homologous (Fig. 6).

**Figure 5.**
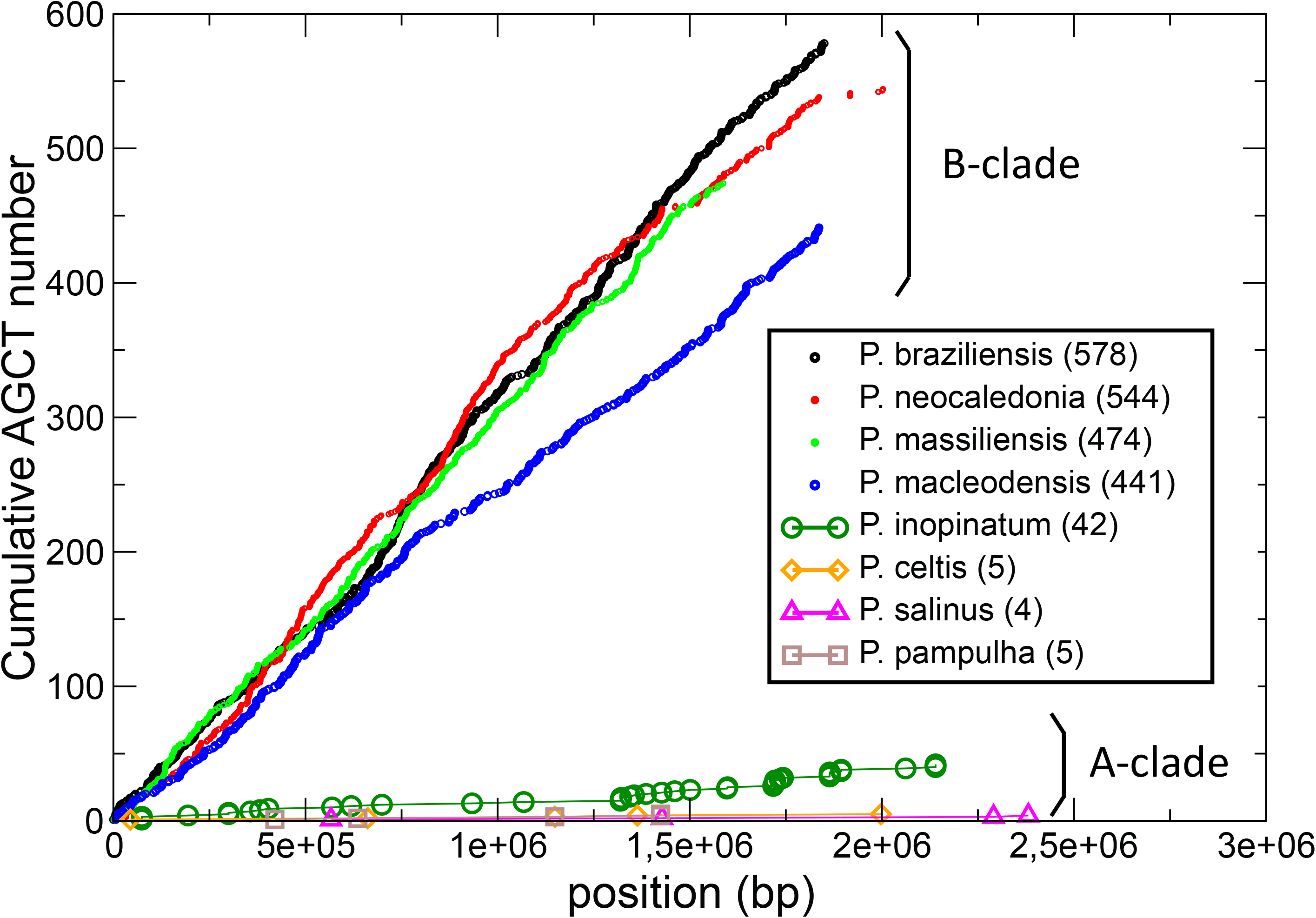
Cumulative distribution of “AGCT” occurrences along the different pandoravirus genomes. The “AGCT” word appears uniformly spread throughout the B-clade pandoravirus genomes, except for a clear rarefaction at the end of the P. neocaledonia genome sequence.

**Figure 6.**
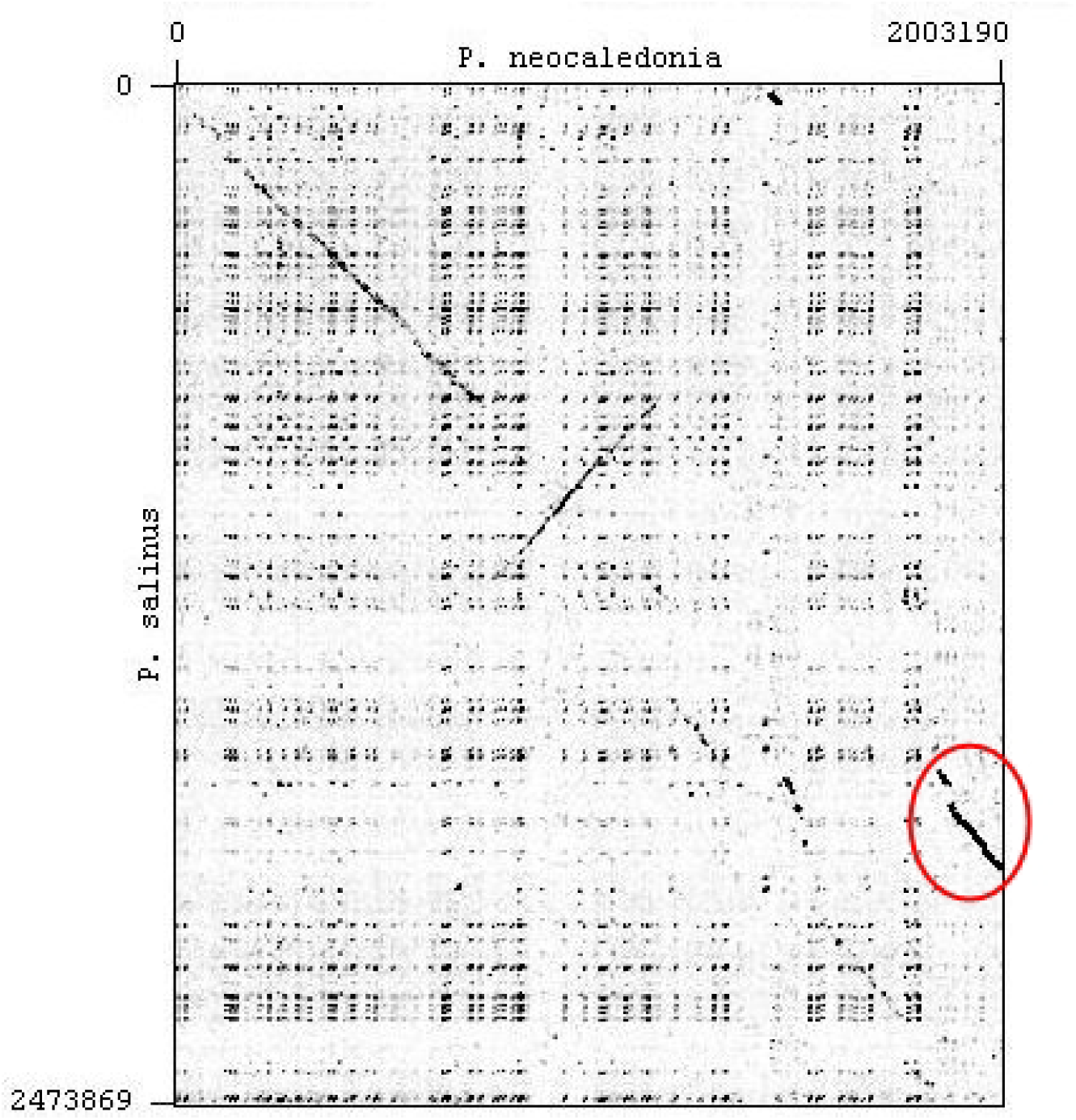
DNA sequence dot-plot comparison of *P. neocaledonia* (horizontal) and *P. salinus* (vertical). The two genomes only exhibit remnants of collinearity except for the terminal region of *P. neocaledonia* (red circle) coinciding with a low “AGCT” density typical of A-clade strains (Fig. 5). Dot plot generated using GEPARD with parameters: word size=15, window size=0 (Krumsiek et al., Bioinformatics **23**, 1026-1028 (2007)).

### “AGCT” was specifically deleted from A-clade pandoravirus genomes

We have seen in the previous section that the extreme difference in the “AGCT” count in *P. dulcis* (N=0) and *P. neocaledonia* (N=544) is not due to the local deletion of an “AGCT”-rich segment. We then investigated if that difference was limited to “AGCT”, or if other 4-mers exhibited large differences in counts. Fig. 7 shows that this was not the case.

**Figure 7.**
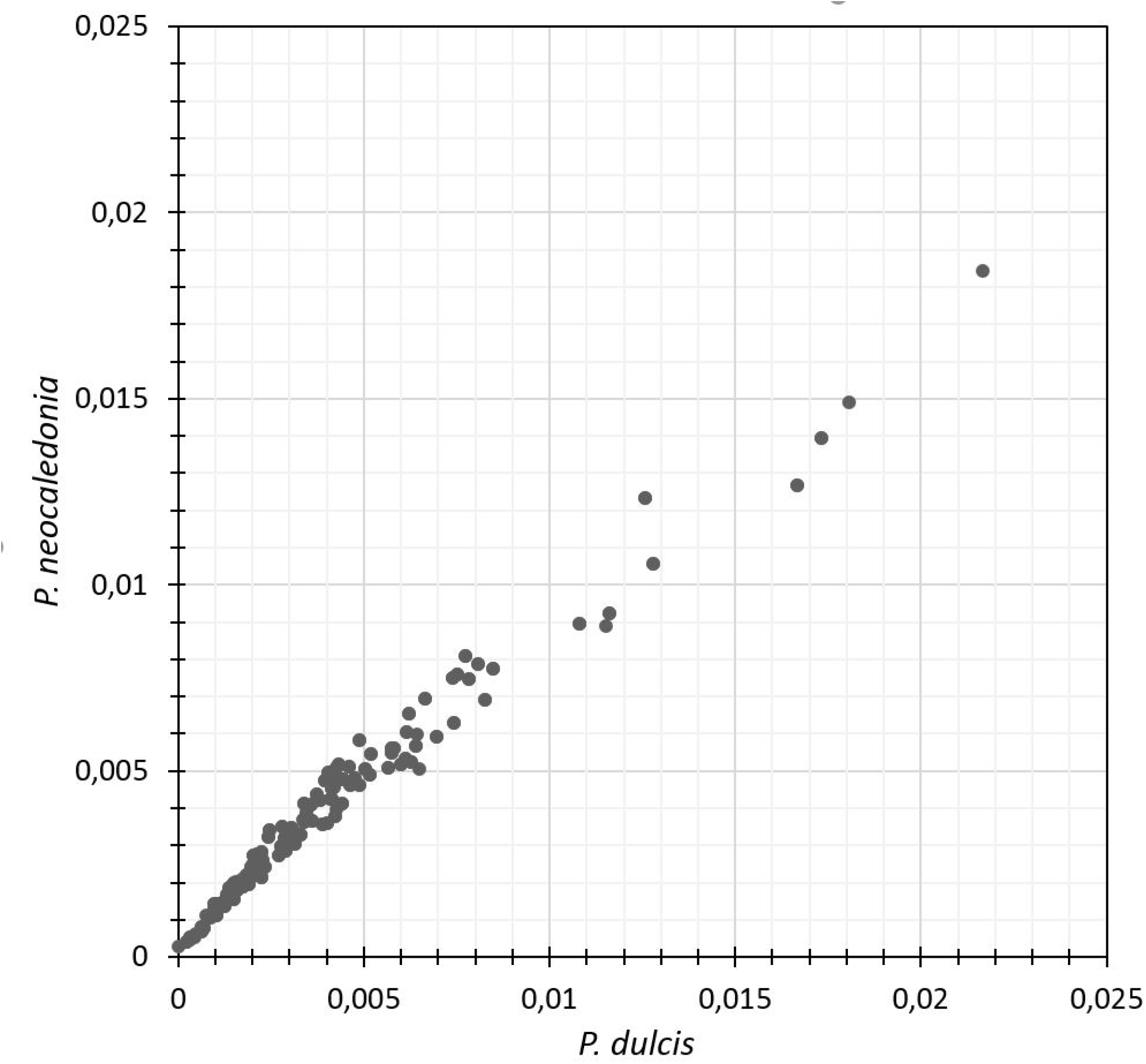
Comparison of the proportion of all 4-mers in *P. dulcis* (A-clade) vs. *P. neocaledonia* (B-clade). The 4 most frequent 4-mers are “GCGC”, “CGCG”, “CGCC”, and “GGCG”.

If the frequencies of the various 4-mers within each genome exhibit tremendous differences (very much at odd with their distribution in randomized sequences, see Fig. 3), the frequency for each 4-mer (low, average or high) was very similar across the two different viral genomes (Spearman correlation, r=0.9859). The difference in “AGCT” count is thus not the consequence of the use of globally distinct 4-mer vocabularies by the two pandoravirus clades. It appears to be due to a selection specifically exerted against the presence of “AGCT” in the genomes of A-clade pandoraviruses.

Another argument in favor of an active selection against the presence of “AGCT” is provided by the following statistical computation. We first identified the orthologous proteins in *P. dulcis* and *P. neocaledonia*, using the best-reciprocal Blastp match criterium. We identified 585 orthologous ORFs. In *P. neocaledonia*, 180 of them were found to contain one or several “AGCT” (for a total of 350 occurrences). We then computed the average percentage of nucleotide identity in the alignments of these 180 *P. neocaledonia* ORFs with their *P. dulcis* orthologous counterparts. The value was 69%.

According to a neutral scenario (and neglecting multiple hits), the probability is thus p = 0.69 that any nucleotide remains the same along the evolutionary trajectory separating the two pandoraviruses. For a given “AGCT”, the probability to remain intact over the same evolutionary distance is *p*_*intact*_ = 0.69^4^ = 0.227, such as none of the four positions is changed. For the sake of simplicity, we will neglect the chance creation of new “AGCT” during the process. As a result, we then expect *P. dulcis* orthologous ORFs to exhibit 68 occurrences (i.e. 0.227 × 350) of “AGCT”.

This simple calculation already indicates that the “AGCT” 4-mer diverged much faster (at least 80 times faster since 350×0.227/80 < 1) than the rest of the orthologous coding regions. This result suggests that the absence of “AGCT” in *P. dulcis* and *P. quercus*, as well as its distinctive low frequency in all A-clade strains is the consequence of an active counter selection. We discuss possible molecular mechanisms in the following section. The above calculation could not be extended to interORFs regions, due to their much lower conservation and their unreliable pairwise alignments.

## Discussion

### Which model for the counter selection of “AGCT”?

Following our statistical computations on random sequences confirmed by the analysis of actual genome sequences, we can safely assume that the genome of the common ancestor of the A- and B-clade pandoraviruses was not missing any 4-mers. Our discussion will thus take for granted that the difference in “AGCT” frequency between the two Pandoraviridae clades is the consequence of a loss in the A-clade rather than a gain in the B-clade.

Any model proposed to explain our results must also take into account that the two types of pandoraviruses are infecting and replicating in the very same Acanthamoeba host. The cause of the marked difference in “AGCT” counts between the two clades (such as a protective mechanism) must thus reside within the viruses themselves. Such inference is further supported by the fact that we found that none of the other families of giant viruses^18^ infecting the very same Acanthamoeba host exhibited a similar 4-mer anomaly in their genome composition.

The first model that comes to mind is inspired from the well-documented restriction-modification systems that many bacteria use to counteract bacteriophage infections. The host bacterial cells express DNA sites (most often short palindromes) specific endonucleases that cut the invading phage genome before it could replicate. Such defense mechanism imposes the bacteria to protect the cognate motif in its own genome using a specific methylase. According to the Red Queen evolutionary concept, the bacteriophages could counteract the host‘s defense by removing the targeted site from their own genome^17^. The absence of the palindrome “GCGC” that we previously noticed in several large enterobacterial phages^13–16^ could result from such evolutionary strategy.

Translating such a model in our system thus requires three distinct assumptions: 1) that the Acanthamoeba cells express an antiviral endonuclease specific for “AGCT”; 2) that B-clade pandoraviruses are immune from it (as other Acanthamoeba-infecting viruses); 3) that A-clade pandoraviruses evolved a different strategy by removing the endonuclease target from their genomes.

Such a model was readily invalidated by simply attempting to digest the B-clade *P. neocaledonia* genomic DNA (extracted from infectious particles) with commercial restriction enzymes (such as PvuII) targeting “cAGCTg” (212 occurrences) and AluI, targeting “AGCT” (544 occurrences). The resulting Pulsed-field gel electrophoresis (PFGE) pattern showed that these sites were not protected (Supplementary Fig.S1). Accordingly, the PacBio data used to sequence the *P. neocaledonia* genome^2^ did not indicate the presence of modified nucleotides at the “AGCT” sites^19^.

We must point out that the above results simultaneously invalidate a symmetrical model where the “AGCT”-specific endonuclease would have been encoded by the pandoraviruses, together with the protective cognate methylase. Such a hijacked restriction/modification system would have been attractive as it is found in chloroviruses^20^, another family of large eukaryotic DNA viruses. Unfortunately, it does not apply here. Accordingly, no homolog of the cognate DNA-methyl transferase was detected among the *P. neocaledonia* or *P. macleodensis* protein-coding gene contents. Further nailing the coffin of such restriction/modification hypothetical model, no difference in terms of potentially relevant endonuclease or DNA methylase was found between the gene contents of the A-clade *P. dulcis* and *P. quercus* and those of the B-clade *P. neocaledonia* and *P. macleodensis*.

A more hypothetical model would assume that the “AGCT” motif is targeted at the transcript level (i.e. “AGCU”) rather than at the DNA level. Classical endonucleases and DNA methylases would thus not be involved in the host-virus confrontation. Several arguments are pleading against a mechanism directly targeting viral transcripts.

First, as B-clade pandoraviruses exhibit similar proportions of “AGCT” in ORFs and inter-ORF regions, the A-clade strains would have had no incentive to eliminate the motif from their intergenic regions, as *P. dulcis* and *P. quercus* have done totally in reaching zero occurrences. “AGCT” is also still present in some protein-coding regions of *P. inopinatum* (N=15), *P. salinus* (N=3), and *P. celtis* (N=1).

Second, very few motif-specific RNAses are known, and to our knowledge, only one is viral: a protein encoded in the bacteriophage T4 RegB gene^21^. We found no significant homolog of this protein in the pandoraviruses or Acanthamoeba. We also looked for mRNA methylases that could act as a protective mechanism for the viral transcript. A single one was described in another family of eukaryotic DNA virus: the product of the Megavirus Mg18 gene^22^. Again, no significant homolog of this protein was detected in the pandoraviruses.

In conclusion to this section, if the presence of “AGCT” is decreasing the virus fitness, we found no evidence that it is due to a DNA or RNA nuclease-mediated defense mechanism. However, it could still be due to an unknown inhibitory mechanism acting at the transcription regulation level to which B-clade pandoviruses would exhibit some immunity. The corresponding proteins could be encoded among the numerous ORFans found in pandoravirus genomes^1–3^.

Finally, could “AGCT” be deleterious for some intrinsic reasons, for instance due to its palindromic structure and composition? This is very unlikely, when one compare the absent “AGCT” in *P. dulcis* and *P. quercus*, with other 4-mers with identical structures and compositions. For instance “ACGT” occurs at 5822 and 6165 positions (in P. dulcis and P quercus, respectively), and “GATC” occurs at 8114 and 8567 times) in (P. dulcis and P. quercus, respectively). The presence or absence of “AGCT” does not either exert a strong constraint on protein sequences, as the amino-acids encoded by “AGC” or “GTC” (Serine and Alanine, respectively) have many possible alternative codons and are easily replaceable residues given their mild physicochemical properties. Finally, we found no evidence that the removal of “AGCT” was due to a specific (for instance, enzyme-mediated) process targeting then replacing the forbidden 4-mer by a constant alternative word. Replacement patterns for 72 *P. dulcis* sites unambiguously mapped to their homologous *P. neocaledonia* “AGCT” counterparts are indicated in supplementary Table S1. It suggests that the complete loss of “AGCT” in the A-clade strains is due to a stringent, nevertheless random (i.e. non-directed) evolutionary process.

The analysis of long nucleotide (and amino acid) sequences as overlapping k-mers has a long history in bioinformatics. Initially proposed in the context of the RNA folding problem^23^, the concept was then quickly applied to many other areas including gene parsing^24^, the detection of regulatory motifs^24, 25^, and has become central to the fast implementation of large-scale similarity search^26, 27^, sequence assembly^28^, and the binning of metagenomics data^29, 30^. However, its popularity should not hide that most of the observed frequency disparities (starting from the simplest mononucleotide composition) between k-mers within a given organism, or across species have not yet received convincing biological explanations^31, 32^. This suggests that profound and unexpected biological insights may one day come out from the analysis of k-mer frequencies, and in particular from their most improbable fluctuations. In a daring parallel with the delayed understanding of the CRISPR/CAS system from the initial spotting of intriguing repeats^33^, we would like to expect that the pandoraviridae “AGCT” distribution anomaly might lead to the discovery of a novel defense mechanism against viral infection.

## Materials and Methods

### Pulse-field gel electrophoresis (PFGE)

Approximately 5,000 pandoravirus particules were embedded in 1% low gelling agarose and the plugs were incubated in lysis buffer (50mM Tris-HCl pH8.0, 50mM EDTA, 1% (v/v) N-laurylsarcosine, 1mM DTT and 1mg/mL proteinase K) for 16h at 50°C. After lysis, the plugs were washed once in sterile water and twice in TE buffer (10mM Tris-HCl pH8.0 and 1mM EDTA) with 1mM PMSF, for15 min at 50°C. The plugs were then equilibrated in the appropriate restriction buffer and digested with 20 units of PvuII or AluI at 37°C for 14 hours. Digested plugs were washed once in sterile water for 15 min, once in lysis buffer for 2h and three times in TE buffer. Electrophoresis was carried out in 0.5X TAE for 18 h at 6V/cm, 120° included angle and 14°C constant temperature in a CHEF-MAPPER system (Bio-Rad) with pulsed times ramped from 0.2s to 120s.

### Availability of data

All virus genome sequences analyzed in this work are freely available from the public GenBank repository (URL://www.ncbi.nlm.nih.gov/genbank/). The Pandoravirus sequences used here correspond to the following accession numbers: *P. dulcis* (NC_021858), *P. neocaledonia* (NC_037666), *P. macleodensis* (NC_037665), *P. salinus* (NC_022098), *P. quercus* (NC_037667), *P. celtis* (NC_), *P. inopinatum* (NC_026440), *P. pampulha* (LT972219.1), *P. massiliensis* (LT972215.1), *P. braziliensis* (LT972217).

---

**Supplementary Figure S1.** Digestion of *P. neocaledonia* DNA at “AGCT” sites. Lane 1: undigested *P. neocaledonia* DNA (2.2 Mb) migrating as expected. The bottom band (below 48.5 kb) correspond to an episome not always present. Lane 2: *P. neocaledonia* DNA digested by the PvuII restriction enzyme (cutting site: cAGCTg). Lane 3: *P. neocaledonia* DNA digested by the AluI restriction enzyme (cutting site: AGCT). These results demonstrate that the “AGCT” sites are not protected by modified nucleotides.

## Acknowledgements

We thank Dr. Sacha Schutz for his inspiring blog (URL: http://dridk.me/) that initiated our interest in the Chaos Game Representation technique. We thank Dr. Matthieu Legendre for verifying the absence of modified nucleotides at “AGCT” sites using the PACBIO sequence data. Our laboratory is supported by the French National Research Agency (ANR-14-CE14-0023-01), France Genomique (ANR-10-INSB-01-01), Institut Français de Bioinformatique (ANR-11-INSB-0013), the Fondation Bettencourt-Schueller (OTP51251), and by the Provence-Alpes-Côte-d’Azur region (2010 12125). We acknowledge the support of the PACA-Bioinfo platform. The funding bodies had no role in the design of the study, analysis, and interpretation of data and in writing the manuscript.

